# Broad spectrum capture of clinical pathogens using engineered Fc-Mannose-Binding Lectin (FcMBL) enhanced by antibiotic treatment

**DOI:** 10.1101/387589

**Authors:** Benjamin T. Seiler, Mark Cartwright, Alexandre L.M. Dinis, Shannon Duffy, Patrick Lombardo, David Cartwright, Elana H. Super, Jacqueline Lanzaro, Kristen Dugas, Michael Super, Donald E. Ingber

## Abstract

FcMBL, an engineered version of the blood opsonin mannose-binding lectin (MBL) that contains the carbohydrate recognition domain (CRD) and flexible neck regions of MBL fused to the Fc portion of human IgG1, has been shown to bind various microbes and pathogen-associated molecular patterns (PAMPs). FcMBL also has been used to create an enzyme-linked lectin sorbent assay (ELLecSA) for use as a rapid (< 1 hr) diagnostic of bloodstream infections. Here we extended this work by using the ELLecSA to test FcMBL’s ability to bind to more than 200 different isolates from over 100 different pathogen species. FcMBL bound to 86% of the isolates and 110 of the 122 (90%) different pathogen species tested, including bacteria, fungi, viruses, and parasites. It also bound to PAMPs including, lipopolysaccharide endotoxin (LPS) and lipoteichoic acid (LTA) from Gram-negative and Gram-positive bacteria, as well as lipoarabinomannan (LAM) and phosphatidylinositol mannoside 6 (PIM_6_) from *Mycobacterium tuberculosis*. The efficiency of pathogen detection and variation between binding of different strains of the same species also could be improved by treating the bacteria with antibiotics prior to FcMBL capture to reveal previously concealed binding sites within the bacterial cell wall. As FcMBL can bind to pathogens and PAMPs in urine as well as blood, its broad-binding capability could be leveraged to develop a variety of clinically relevant technologies, including infectious disease diagnostics, therapeutics, and vaccines.

## Introduction

Mannose-binding lectin (MBL) is a key host-defense protein associated with the lectin pathway of the innate immune system [1], and deficiency of MBL can lead to increased susceptibility to a wide-spectrum of infectious diseases [2–4]. MBL functions as a calcium-dependent, pattern-recognition opsonin that binds a range of carbohydrate molecules associated with the surfaces or cell walls of many different types of pathogens [5]. Collectively these microbial surface carbohydrate molecules, including for example, lipoteichoic acid (LTA) and lipopolysaccharide endotoxin (LPS), are referred to as pathogen-associated molecular patterns (PAMPs) [6,7]. MBL has the intrinsic ability to distinguish foreign PAMPs from self, subsequently activating the complement system and providing protection via antibody-dependent and independent mechanisms [8,9].

Due to the evolutionary conserved recognition carbohydrate moieties of PAMPs, MBL is a broad spectrum opsonin that can bind over 90 different species of pathogens, including Gram-negative and Gram-positive bacteria, fungi, viruses, and parasites [10–14]. MBL binding to these various pathogens has been demonstrated by means of flow cytometry [14,15], radio-immunoassay [13,16], enzyme-linked immunosorbent assay (ELISA) [13,17], immunofluorescence and scanning electron microscopy (SEM) [18], and *Saccharomyces cerevisiae*-induced MBL activation and bystander lysis of chicken erythrocytes [19]. However, many discrepancies in MBL binding have been described, depending on the method used. For example, use of flow cytometry revealed little to no MBL binding to *Pseudomonas aeruginosa*, while others have reported good binding of MBL to *Pseudomonas aeruginosa* using a hemolytic assay [15,19].

We set out to address these conflicting results by leveraging recent development of an engineered version of MBL that contains the carbohydrate recognition domain (CRD) and flexible neck regions of MBL fused to the Fc portion of human IgG1, which is known as FcMBL [20]. The engineered FcMBL lacks the regions of the native molecule that interact with MBL-associated serine proteases (MASPs) that activate complement and promote blood coagulation, and thus, it can be used to capture of PAMPs from complex biological fluids, such as blood and urine, without activating effector functions of complement, coagulation, and phagocytosis. We have previously used FcMBL in extracorporeal therapies, such as hemofiltration, and in diagnostics to capture and detect *Staphylococcus aureus* from osteoarticular and synovial fluids of infected patients [20–22]. In the present study, we used a previously described sandwich enzyme-linked lectin sorbent assay (ELLecSA) in which both live and fragmented pathogens (PAMPs) are captured magnetically using FcMBL conjugated to magnetic beads and then detected with horseradish peroxidase (HRP)-labeled MBL [23]. This ELLecSA has enabled rapid (< 1 hr) diagnosis of bloodstream infections by capturing and detecting PAMPs in whole blood from human patients [23]. Here we use this ELLecSA to measure direct binding of FcMBL to over 200 pathogen isolates from over 100 different pathogen species, including bacteria, fungi, viruses, and parasites, as well as bacterial cell wall antigens. We demonstrate that FcMBL binds 90% of the pathogen species tested and that antibiotic treatment of the bacterial pathogens exposes previously concealed FcMBL binding sites on cell walls, thus increasing the efficiency of pathogen detection and reducing variation between binding of different strains of the same species. We also show FcMBL can detect PAMPs in urine as well as blood, making this potential diagnostic technology highly synergistic with standard of care antibiotic therapy.

## Results

### FcMBL binding to bacteria

We first set out to determine the range of pathogens that FcMBL can capture by screening multiple species of bacteria, fungi, viruses, and parasites using the ELLecSA detection technology. In the FcMBL ELLecSA, pathogen materials in experimental samples are captured with FcMBL immobilized on superparamagnetic beads (1 μm diameter), magnetically separated, washed, bound to human MBL linked to horseradish peroxidase (HRP), magnetically separated again, washed, and then tetramethylbenzidine (TMB) substrate is added to quantify the amount of pathogen material bound (**Fig 1**). Initially our focus was on screening bacteria and as such we compiled a comprehensive list of clinically relevant bacterial pathogens (**Table 1**). When we screened 88 different species of bacteria, we found that FcMBL detected 69 out of 88 live microbes (78%) and that more could be detected (76 out of 88; 86%) after they were treated with antibiotics (**Table 1**). The antibiotics we used in this study were clinical grade cefepime, ceftriaxone, meropenem, amikacin, and vancomycin, to provide enough coverage to target this diverse range of bacteria. We dosed each bacterial class with a single appropriate antibiotic at a dose (1 mg/mL) much higher than their minimal inhibitory concentrations (MIC) to obtain acute fragmentation within 4 hours. Pseudomonas was treated with the 4^th^ generation cephalosporin, cefepime, due to weak coverage by 3^rd^ generation ceftriaxone, and methicillin-resistant *Staphylococcus aureus* (MRSA) was treated with vancomycin.

**Fig 1.**
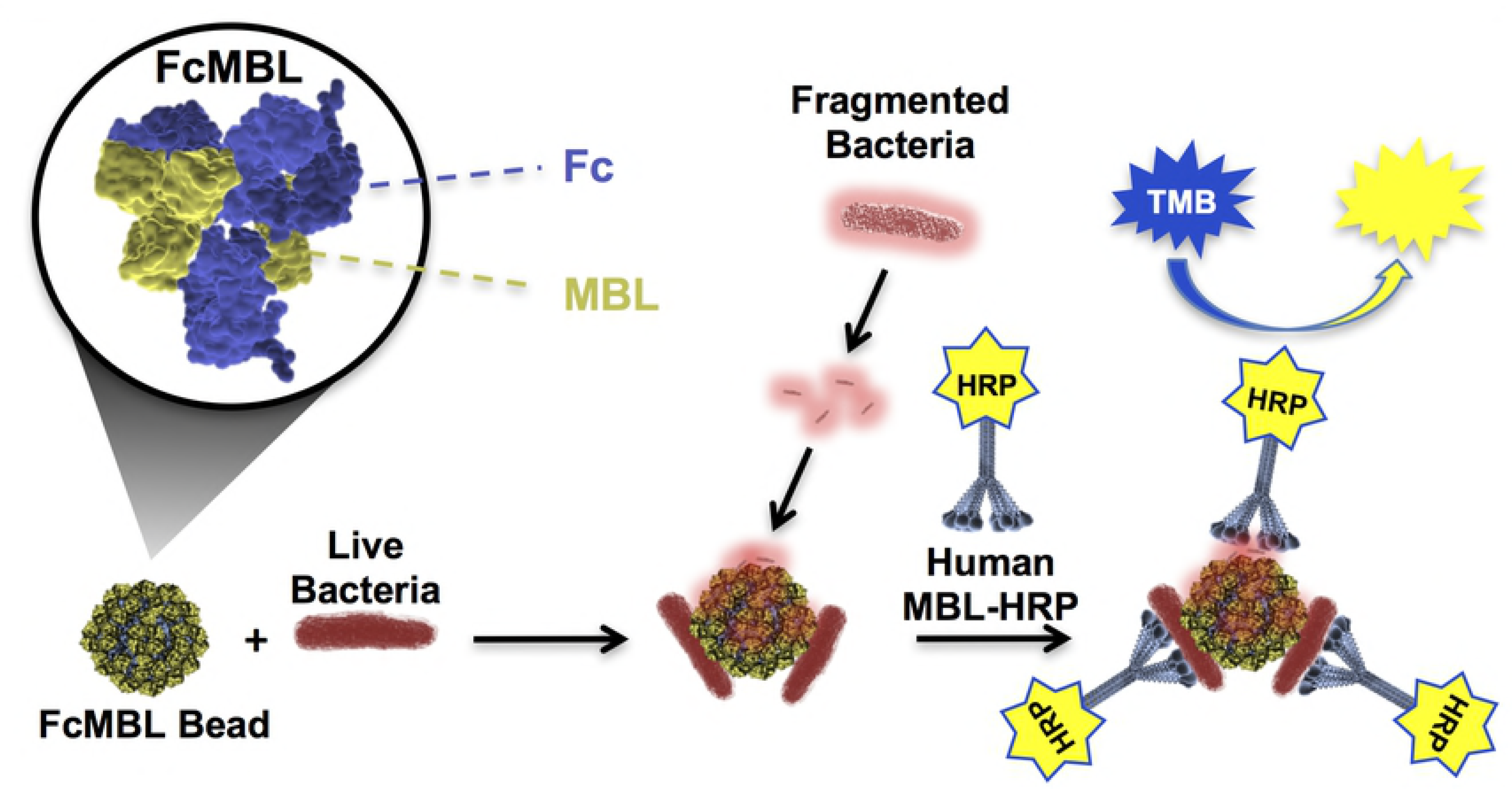
Diagrammatic representation of the FcMBL ELLecSA. An N-terminal aminooxy-biotin on the Fc allows oriented attachment to streptavidin coated superparamagnetic beads (FcMBL Bead). FcMBL beads capture live and fragmented bacteria, which are then magnetically separated and detected using recombinant human MBL linked to horseradish peroxidase (Human MBL-HRP). Tetramethylbenzidine (TMB) substrate is added to quantify captured bacteria and results are read at OD 450nm.

**Table 1.**
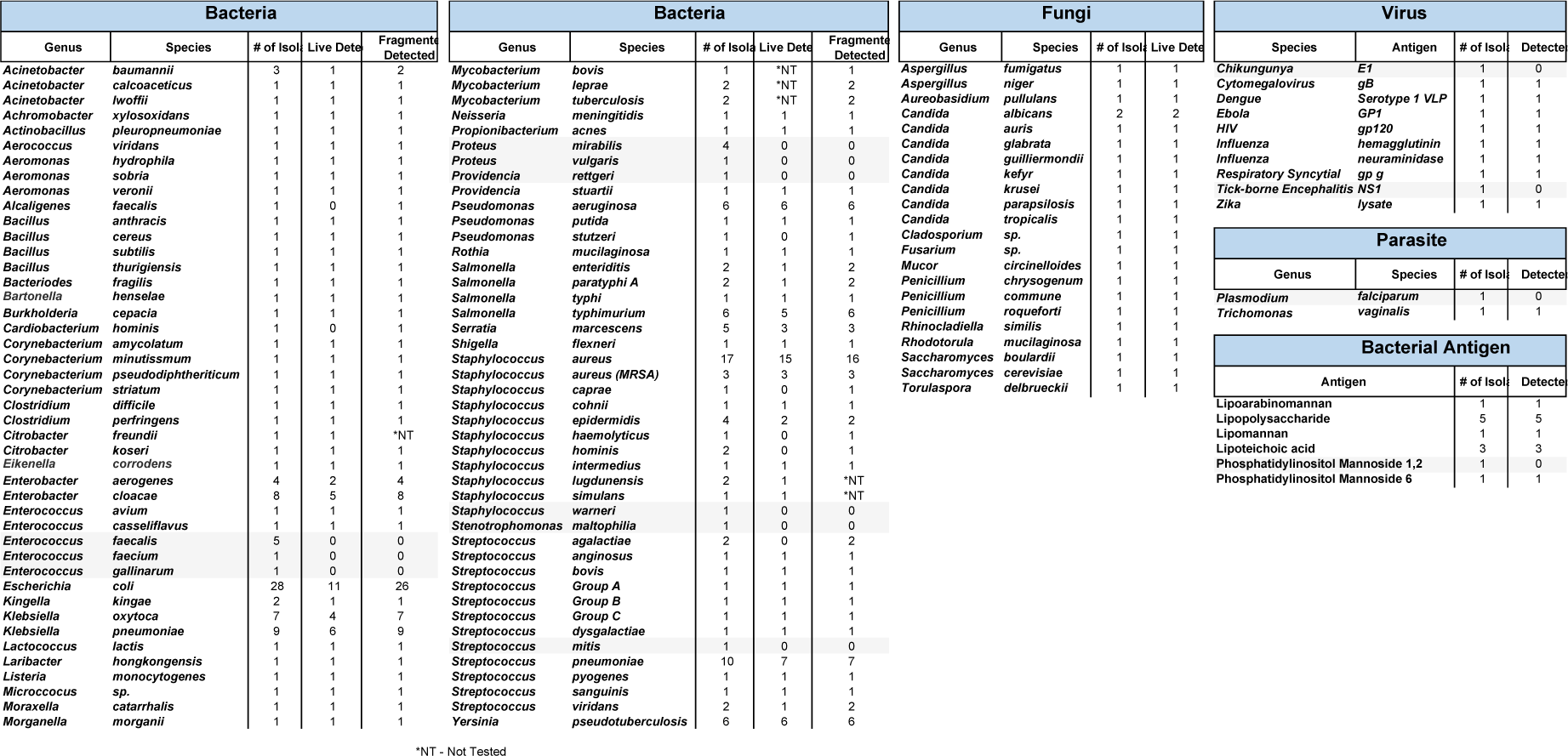
FcMBL binding profile of bacteria, fungi, viruses, parasites, and bacterial antigens determined by FcMBL ELLecSA.

Multiple species of bacteria, including multiple isolates (# of isolates), were screened to determine FcMBL binding. Total number detected of both live and fragmented bacterial isolates is shown. Fungi were screened and total number detected for live isolates shown. Purified or inactivated viral, parasite, and bacterial antigens were tested directly in TBST 5mM CaCl_2_ buffer, and number detected shown. Test samples were performed in duplicate. *NT indicates not tested.

To determine if inducing bacterial fragmentation via antibiotic treatment would produce different effects on FcMBL detection sensitivity when tested using different strains of the same species, we screened 137 isolates from 22 of the 88 Gram-positive and Gram-negative bacterial species, including antibiotic-resistant organisms (e.g., MRSA) (**Fig 2**). As before, FcMBL bound a greater proportion of the pathogens when fragmented with antibiotic treatment (115/137 = 84%) than when live and intact (80/137 = 58%). For some bacterial species such as *Enterobacter cloacae*, *Escherichia coli*, *Klebsiella oxytoca*, and *Klebsiella pneumoniae* we found antibiotic-induced fragmentation greatly increased FcMBL binding, whereas other bacteria like *Pseudomonas aeruginosa*, *Yersinia pseudotuberculosis*, and MRSA bound equally well when live and intact (**Fig 2**). With the exception of *Proteus mirabilis* and *Enterococcus faecalis*, which FcMBL did not bind at all, the capture of fragmented bacteria was equal to or greater than that of live bacteria.

**Fig 2.**
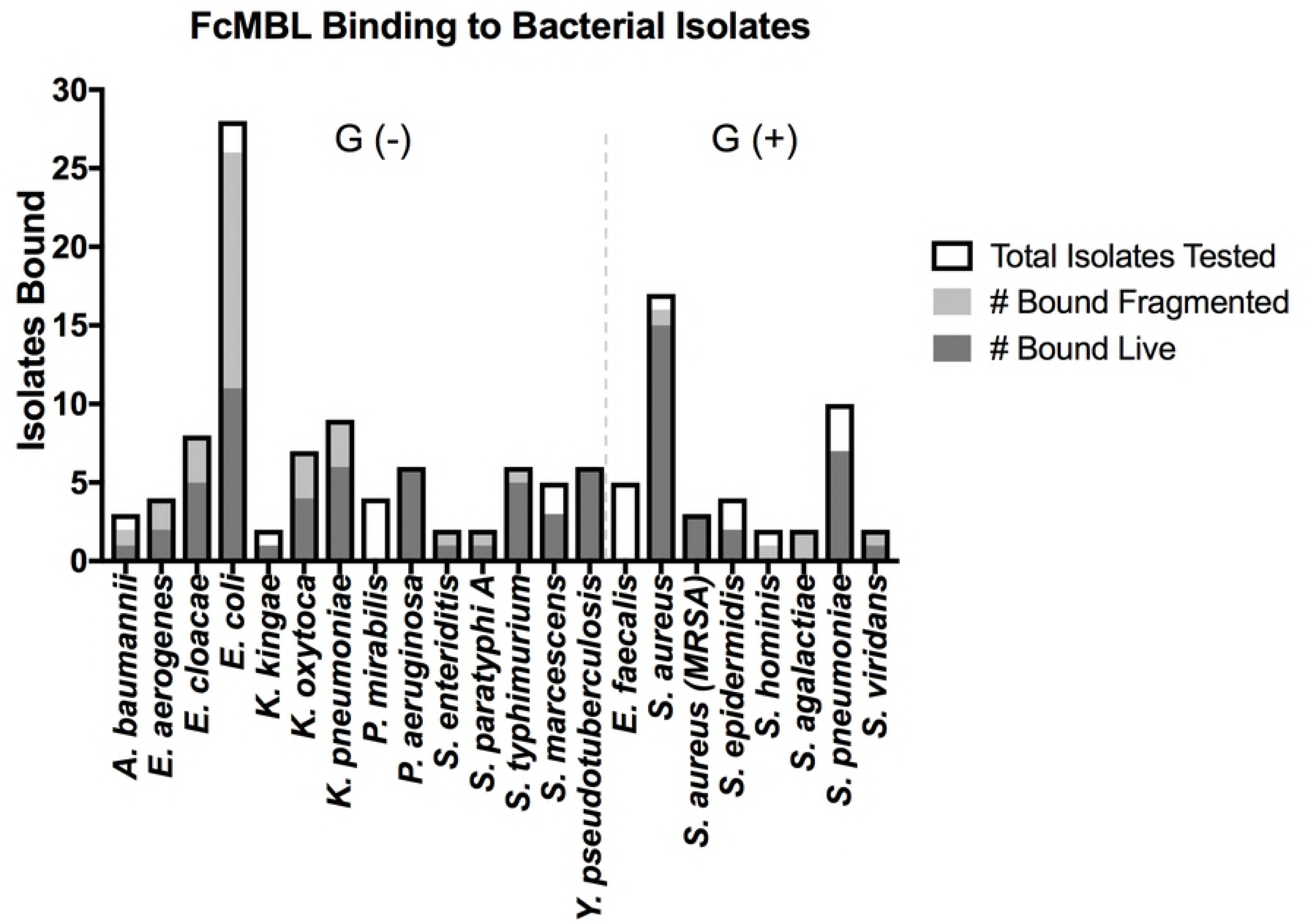
FcMBL binding to bacterial isolates is equal to or enhanced with antibiotic fragmentation. Graph is divided between Gram-negative bacterial isolates [Gram (-)] and Gram-positive bacterial isolates [Gram (+)]. Data are presented as the number of bacterial isolates bound live and fragmented within the total isolates tested for each species: *A. baumannii* (*n* = 3), *E. aerogenes* (*n* = 4), *E. cloacae* (*n* = 8), *E. coli* (*n* = 28), *K. kingae* (*n* = 2), *K. oxytoca* (*n* = 7), *K. pneumoniae* (*n* = 9), *P. mirabilis* (*n* = 4), *P. aeruginosa* (*n* = 6), *S. enteriditis* (*n* = 2), *S. paratyphi A* (*n* = 2), *S. typhimurium* (*n* = 6), *S. marcescens* (*n* = 5), *Y. pseudotuberculosis* (*n* = 6), *E. faecalis* (*n* = 5), *S. aureus* (*n* = 17), *S. aureus*(MRSA) (*n* = 3), *S. epidermidis* (*n* = 4), *S. hominis* (*n* = 2), *S. agalactiae* (*n* = 2), *S. pneumoniae* (*n* = 10), *S. viridans* (*n* = 2).

These findings are consistent with past studies that showed the efficiency of MBL binding to live bacteria differs between isolates from the same bacterial genus and species, possibly due to differences in encapsulation [15,16]. Here we demonstrate this heterogeneity between MBL binding live isolates of the same species, but show upon exposure of previously cryptic binding sites using antibiotic disruption, that we were able to bind FcMBL to both isolates of the same species. To illustrate this point, we show the clinical isolates, *Escherichia coli* 41949 and *Streptococcus pneumoniae* 3, exhibited equivalent FcMBL binding whether they were live or fragmented with antibiotics (1 mg/mL cefepime or ceftriaxone, respectively, for 4 hours), whereas fragmented forms of *E. coli* RS218 and S*. pneumoniae* 19A isolates bound much more effectively to FcMBL than living forms (**Fig 3A-D**). This difference was further supported visually using scanning electron microscopy (SEM) in which magnetic FcMBL beads could be seen to bind both live and fragmented versions of *E. coli* 41949 and *S. pneumoniae* 3, but with *E. coli* RS218 and *S. pneumoniae* 19A, the FcMBL beads only bound to fragmented material (**Fig 3E-H**). FcMBL binding due to increases in fragmentation also correlated with LPS release measured using a limulus amebocyte lysate (LAL) assay: equal amounts of LPS were detected for *E. coli* 41949 whether live or fragmented, whereas LPS levels were higher in antibiotic-treated *E. coli* RS218 (**Fig 3I,J**). These results suggest that antibiotic treatment results in exposure of previously cryptic PAMPs in the cell wall, including toxins such as LPS, which leads to greatly increased binding of FcMBL.

**Fig 3.**
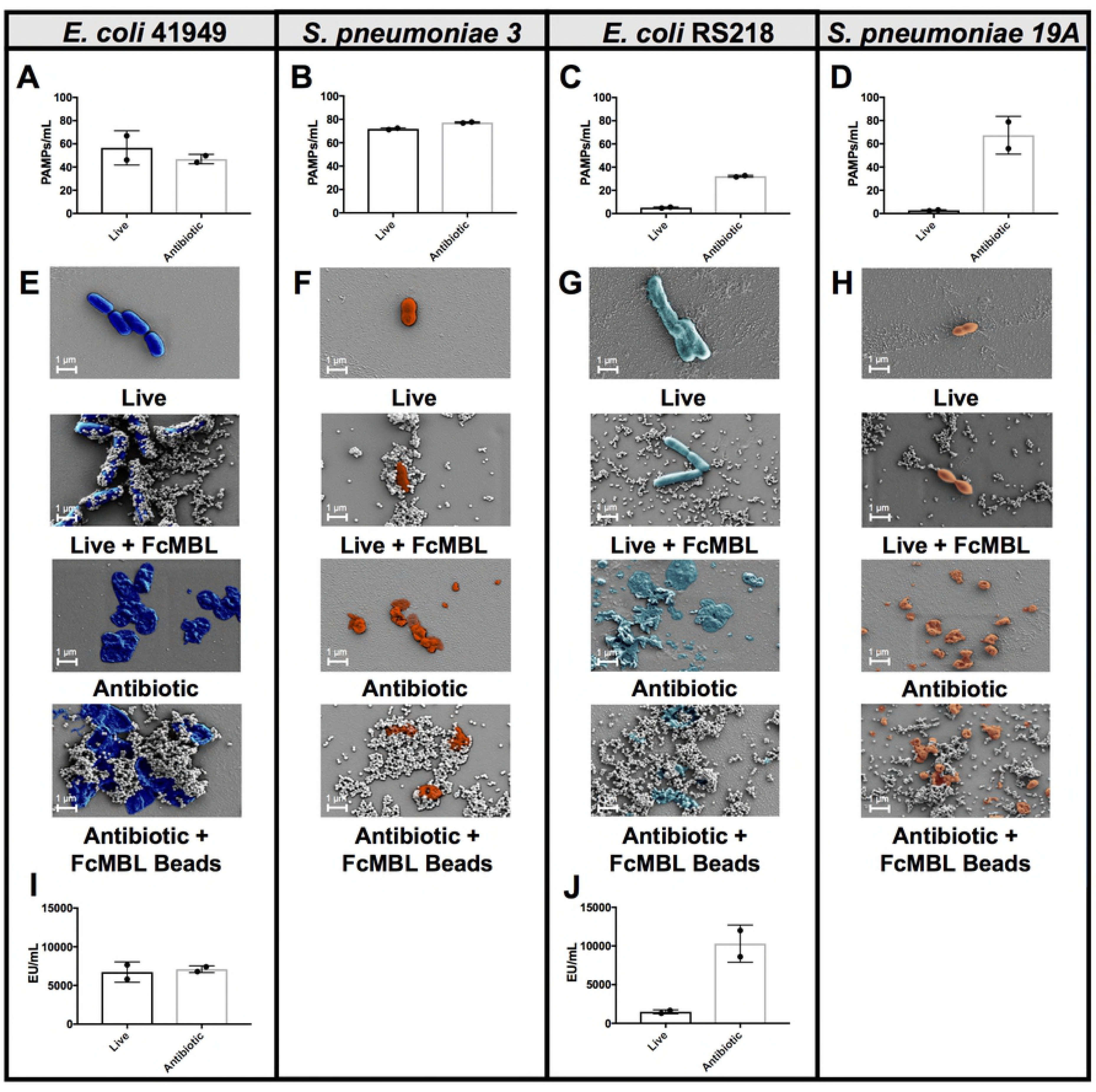
FcMBL bacterial binding efficiency can be enhanced with antibiotic treatment. **(A-D)** PAMPs/mL detection by FcMBL ELLecSA of both live (1e7 CFU/mL) and fragmented bacteria using antibiotics (cefepime 1mg/mL or ceftriaxone 1 mg/mL). **(A)** *E. coli* 41949, **(B)** *S. pneumoniae* 3, **(C)** *E. coli* RS218, and **(D)** *S. pneumoniae* 19A. **(E-H)** Scanning electron microscopy images showing FcMBL bead (128 nm) capture of both live and fragmented bacteria using antibiotics (cefepime 1mg/mL or ceftriaxone 1 mg/mL). **(E)** *E. coli* 41949, **(F)** *S. pneumoniae* 3, **(G)** *E. coli* RS218, and **(H)** *S. pneumoniae* 19A. **(I, J)** LPS endotoxin measurement (LAL assay) using 1e7 CFU/mL of both live and fragmented bacteria using antibiotics (cefepime 1 mg/mL). **(I)** *E. coli* 41949 and **(J)** *E. coli* RS218.

In these studies, we found that 9 bacterial species, including multiple species of enterococcus and proteus, failed to bind to FcMBL even when treated for 4 hours with combinations of antibiotics (500 µg/mL vancomycin and 500 µg/mL amikacin for Gram-positive isolates or 500 µg/mL cefepime + 500 µg/mL amikacin for Gram-negative isolates) (**Fig 2** and **Table 1**). Importantly however, FcMBL was able to detect 85% of the bacterial isolates (**Fig 4**), which includes 9 of the 10 pathogens responsible for most healthcare-associated infections in acute care hospitals in the U.S., with enterococcus species being the one exception [24].

**Fig 4.**
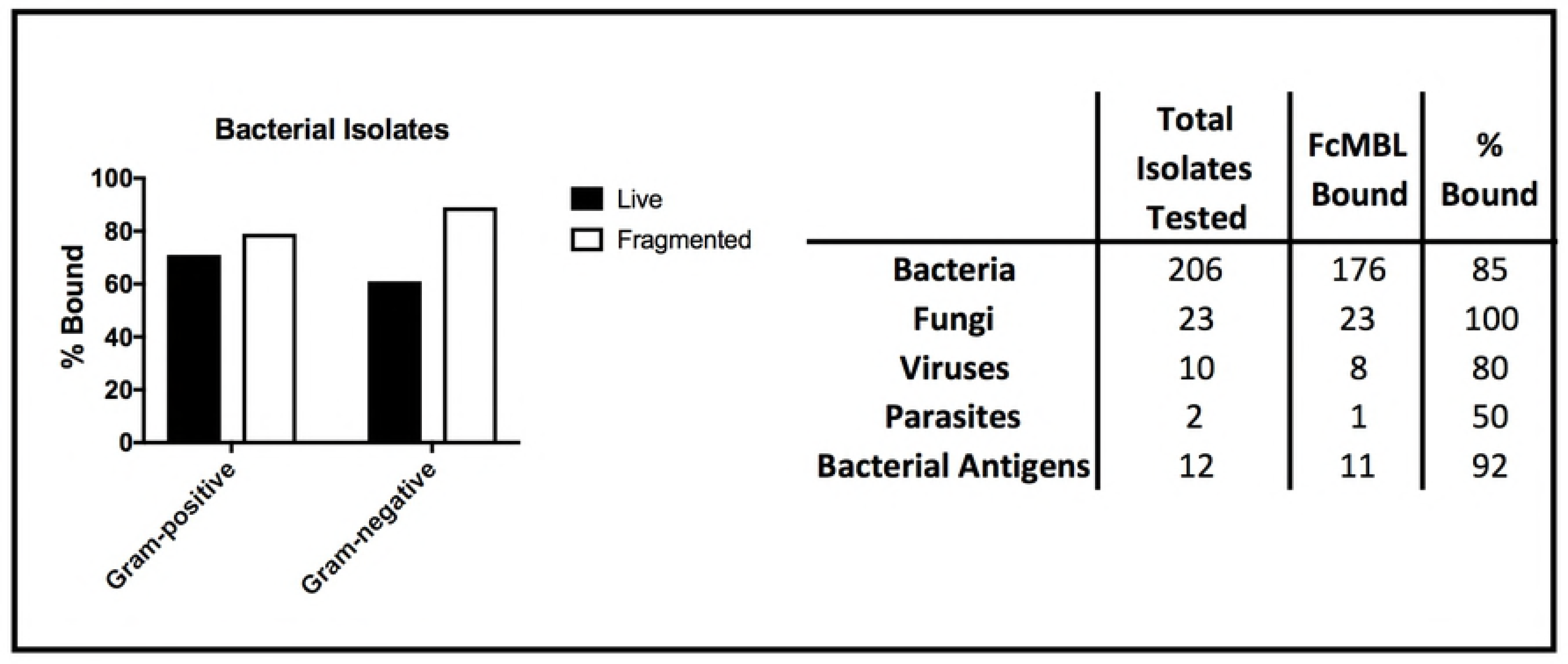
Summary of FcMBL pathogen capture. **(Left)** Percent of live and fragmented Gram-negative (*n* = 119) and Gram-positive (*n* = 78) bacterial isolates bound by FcMBL ELLecSA. **(Right)** Chart showing total number of isolates tested by FcMBL ELLecSA for bacteria, fungi, viruses, parasites, and bacterial antigens, total number FcMBL bound, and total percent bound overall.

### FcMBL binding to bacterial cell wall components

We further explored FcMBL’s ability to bind cell wall components because when antibiotics were used to disrupt the membranes of Gram-negative isolates (*n* = 119), there was a significant boost in FcMBL detection efficiency with fragmented cells (89%) versus live intact cells (61%) (**Fig 4**), and this is likely due to exposure of LPS that is present in high concentrations in their cell wall [25]. In contrast, FcMBL detected a greater percentage (71%) of live Gram-positive isolates (*n* = 78), and this only slightly increased to 79% after antibiotic treatment (**Fig 4**) Thus, to better understand some of the major targets that FcMBL binds when bacteria are fragmented, we extended our analysis using purified samples of the hallmark PAMPs, LPS and LTA [25,26].

Using the ELLecSA, we screened LPS purified from Gram-negative bacteria (*Serratia marcescens, Klebsiella pneumoniae*, and *Salmonella enterica* serovar enteritidis), as well as LTA from Gram-positive bacteria (*Enterococcus hirae*, *Staphylococcus aureus*, and *Streptococcus pyogenes)*. The ability of FcMBL to target these PAMPs was quantified in buffer (50mM Tris-HCl, 150mM NaCl, 0.05% Tween-20, pH 7.4 supplemented with 5mM CaCl_2_ [TBST 5mM CaCl_2_]) to promote optimal MBL calcium-dependent binding, as well as in more clinically relevant human whole blood samples. We found that FcMBL was able to detect LPS from all 3 Gram-negative species in blood, however, the sensitivity was consistently lower than that detected in buffer (**Fig 5A-C**). FcMBL also bound to *E. hirae* LTA very well (15.6 ng/mL limit of detect in buffer and 62.5 ng/mL in blood) (**Fig 5D**), which is consistent with past findings [27,28]. Interestingly, FcMBL also bound to LTA from both *S. aureus* (**Fig 5E**) and *S. pyogenes* (**Fig 5F**) even though the same report that described MBL binds to *E. hirae* LTA claimed that it does not bind LTA from these species due to lack of glycosyl substituents [27,28].

**Fig 5.**
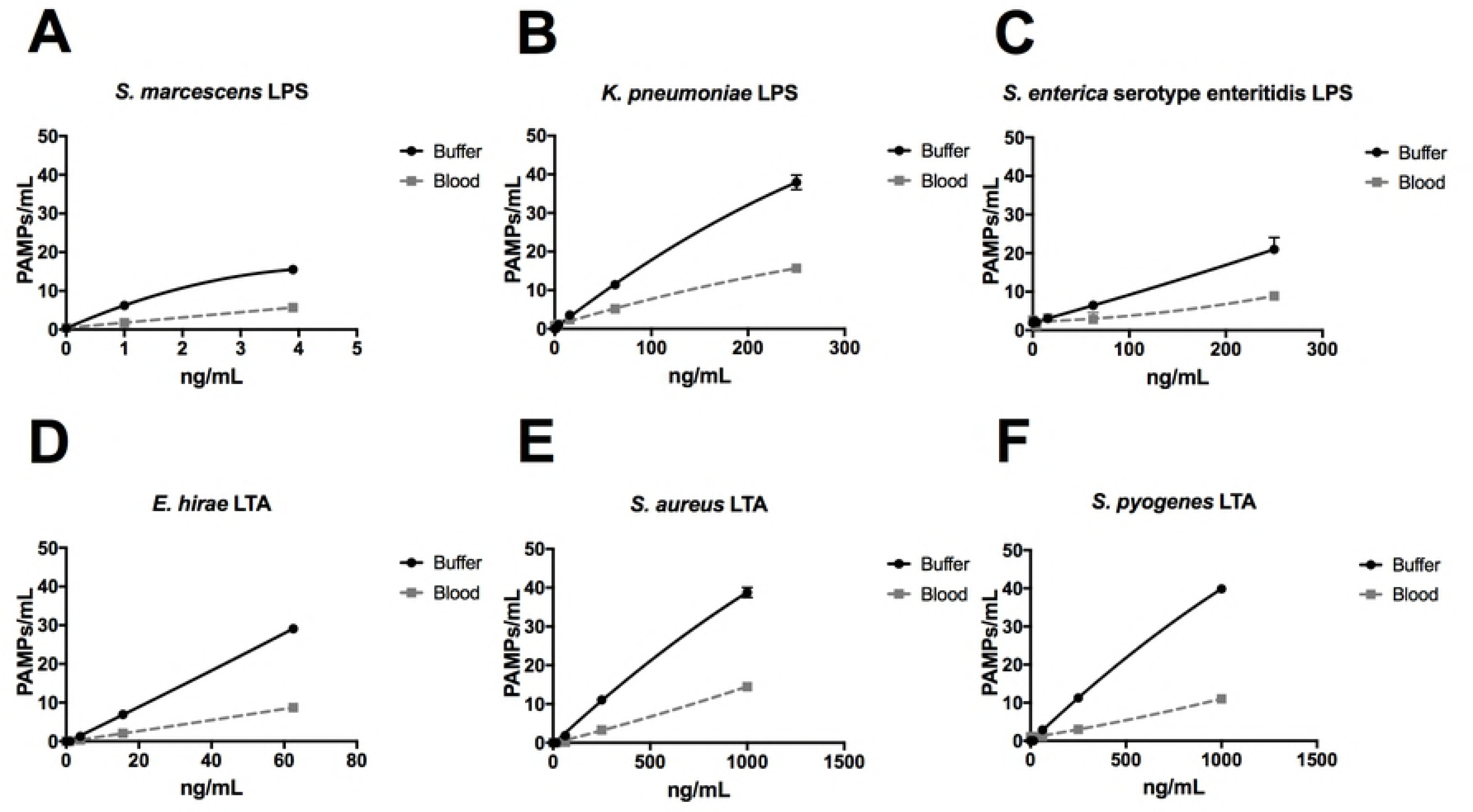
FcMBL ELLecSA screening of purified lipopolysaccharide (LPS) and lipoteichoic acid (LTA). LPS from **(A)** *S. marcescens*, **(B)** *K. pneumoniae*, and **(C)** *S. enterica* serovar enteritidis, and LTA from **(D)** *E. hirae*, **(E)** *S. aureus*, and **(F)** *S. pyogenes* were spiked into either TBST 5mM CaCl_2_ buffer or whole human blood at indicated concentrations.

We next tested FcMBL’s ability to bind lipoarabinomannan (LAM) and its biosynthetic precursors, phosphatidylinositol mannoside 1 & 2 and 6 (PIM_1,2_ and PIM_6_) from *Mycobacterium tuberculosis* (TB) strain H37Rv [29,30]. LAM released from metabolically replicating or degrading TB bacteria has been detected in both blood and urine [31,32]. Thus, we assessed the ability of FcMBL to capture and detect LAM, as well as PIM_1,2_ and PIM_6_, spiked into both of these complex biological fluids as well as buffer. Our initial screen in buffer confirmed that FcMBL can detect LAM and PIM_6_ at levels down to 1 ng/mL, but it did not detect PIM_1,2_ (**Fig 6A-C**). FcMBL also bound to LAM in both blood and urine but its binding sensitivity was reduced as it could only detect 15.6 ng/mL. FcMBL binding to PIM_6_ exhibited a similar sensitivity in buffer, but it could only detect 62.5 ng/mL and 4 ng/mL in blood and urine, respectively.

**Fig 6.**
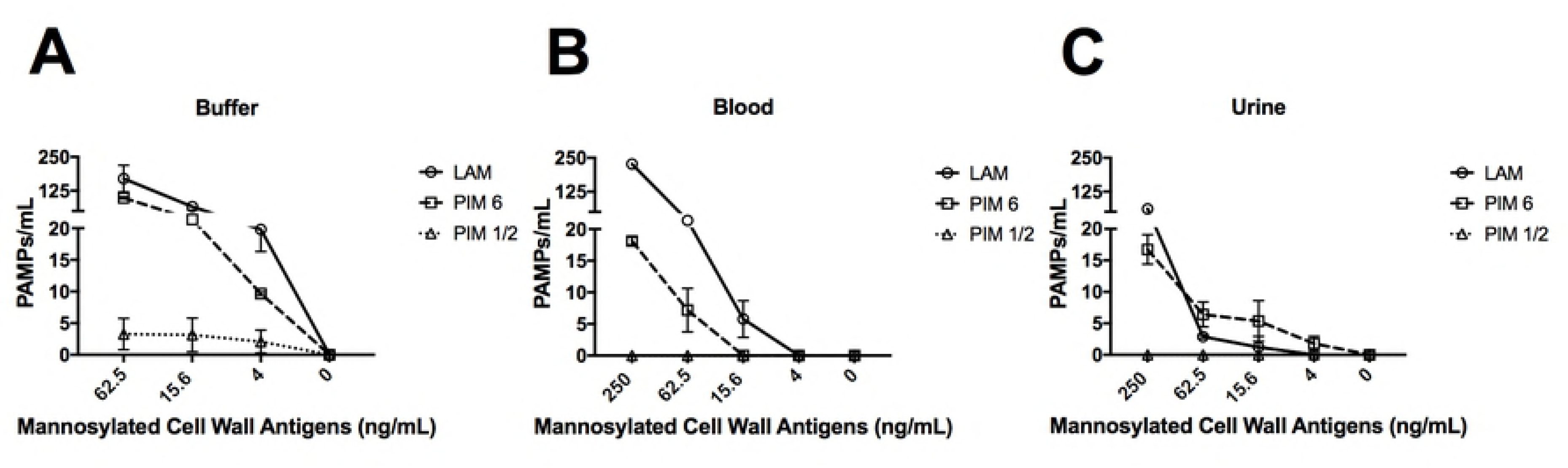
FcMBL ELLecSA screening of *Mycobacterium tuberculosis* strain H37Rv glycolipids PIM_1,2_, PIM_6_, and LAM. Glycolipids spiked into **(A)** buffer (TBST 5mM CaCl_2_), **(B)** whole human blood, and **(C)** urine. FcMBL detected PIM_6_ and LAM at 1 ng/mL in buffer, but not PIM_1,2_. Sensitivity of PIM_6_ and LAM is reduced in whole human blood and urine.

### FcMBL binding to fungi, parasites, viruses, and bacterial cell wall antigens

In addition to screening multiple bacteria, we also tested FcMBL’s ability to bind to 22 different species of fungi, 10 species of virus, 2 species of parasites, and 6 types of purified bacterial cell wall antigens (**Table 1**). In contrast to studies with bacteria, FcMBL was found to bind 100% of live fungal cells from all 22 species and 23 isolates tested (**Fig 4**). Of the two parasites tested in this preliminary analysis, only *Trichomonas vaginalis* was bound by FcMBL, whereas 80% of the viruses screened and 92% of the purified bacterial cell wall antigens were detected (**Fig 4**). The handful of pathogen material FcMBL did not detect included the E1 protein from chikungunya virus, the NS1 protein from tick-borne encephalitis virus, *Plasmodium falciparum*, and PIM_1,2_ from TB. In total, the overall FcMBL binding profile respectively detected 208 (86%) of the 241 isolates and 110 (90%) of the 122 different pathogen species tested.

## Discussion

MBL has been reported to bind to over 90 different pathogen species as well as PAMPs released from these microbes based on studies in which binding was assessed by means of flow cytometry, ELISA, radio-immunoassay, immunofluorescence and SEM, or hemolytic assays [12–19]; however, different results have been obtained with different methods. Here we explored the broad spectrum binding capabilities of an engineered form of MBL, known as FcMBL, using a previously described magnetic ELLecSA detection assay to quantify binding of MBL to over 200 different bacteria, fungi, viruses, parasites, and bacterial cell wall antigens. FcMBL was previously shown to bind to PAMPs released from 47 of 55 (85%) microbial species tested, including 38 species of bacteria and 9 species of fungi [23]. The FcMBL ELLecSA also was able to detect infectious PAMPs in whole blood of sepsis patients, regardless of antibiotic therapy (blood culture positive or negative) with a detection sensitivity and specificity of 85% and 89%, respectively [23]. In the present study, we utilized the ELLecSA to compile a more comprehensive pathogen binding profile composed of over 200 isolates from more than 100 different pathogen species, which include not only bacteria and fungi, but also viruses, parasites, and bacterial cell wall antigens. Our results confirm that FcMBL binds to 86% of the isolates and 110 of the 122 species tested, which corresponds to a 90% detection sensitivity.

MBL binding to different clinical bacterial isolates of the same species has previously produced conflicting results [15,16]. These same studies also described that most Gram-negative isolates (encapsulated strains) bound little or no MBL. We have reported similar results as we previously found that FcMBL only bound 38% of live clinical *E. coli* isolates tested; however, upon fragmentation and release of PAMPs, FcMBL detection of these same isolates increased to 92% [23]. Broader examination of Gram-negative bacteria in the present study revealed a similar pattern: FcMBL only detected 73/119 (61%) of live isolates, but when these same microbes were treated with antibiotics, the detection sensitivity increased to 89% (106/119 isolates). Apparently, by treating the bacteria with antibiotics, we were able to disrupt the encapsulated cell wall, exposing and presenting previously hidden PAMPs, thereby increasing binding and reducing variability between isolates within the same bacterial species. However, even with cell wall disruption, FcMBL did not bind 9 bacterial species, including multiple isolates of *E. faecalis* and *P. mirabilis*. These microbes likely lack the complex polysaccharide antigens which FcMBL and MBL bind. Alternatively, the binding sites might still be present, but if they are, they remain inaccessible due to the unique structure of their cell wall (e.g. carbohydrate conformation, sugar density or composition). Alternatively, the antibiotics we used might not be optimal for disrupting the cell wall in these cells.

To emphasize the ability of FcMBL to be used to detect the presence of a systemic pathogenic infection even when blood cultures are negative, we tested its ability to bind LPS and LTA that are major PAMP-associated toxins released by multiple species of bacteria. FcMBL was able to detect both LPS and LTA from all 6 bacterial species tested in both buffer and blood, although detection sensitivity was consistently higher in buffer. In addition, we explored whether FcMBL binds to the antigenic PAMPs, LAM, PIM_1,2_, and PIM_6_ from *M. tuberculosis* H37Rv because these are active virulence factors associated with TB pathogenesis, and hence, they are critically important targets for point-of-care diagnostic and vaccine applications [33–36]. We found that FcMBL can detect LAM and PIM_6_, but not PIM_1,2_, in buffer, urine, and blood; this difference in binding is likely due to the fact that PIM_1,2_ has 4 fewer branched mannose residues than PIM_6_ [37].

In summary, FcMBL’s ability to both bind to numerous types of infectious pathogens and capture many of the cell wall PAMPs released by these microbes when treated by antibiotics, in complex biological fluids further demonstrates the potential value of using FcMBL capture for rapid detection of bloodstream infections, even when blood cultures are negative. To our knowledge, this is the broadest range and largest number of pathogens and PAMPs that have been shown can be detected by a single blood opsonin or lectin. FcMBL’s ability to detect cell wall fragments synergizes well with standard of care antibiotic therapy, and it’s broad-range pathogen capture and detection can be leveraged to develop a wide range of infectious disease diagnostics, therapeutics, and vaccines.

## Materials and methods

### Pathogen sources

Bacteria, fungi, viruses, parasites, and bacterial cell wall antigens were obtained from a multitude of sources which include: Abcam (Cambridge, USA), AERAS (Rockville, USA), American Type Culture Collection (Manassas, USA), Biodefense and Emerging Infections Resources (Manassas, USA), Boston Children’s Hospital (Boston, USA), Brigham and Women’s Hospital Crimson Biorepository (Boston, USA), Hospital Joseph-Ducuing (Toulouse, France), Sigma-Aldrich (St. Louis, USA), Sino Biological (Beijing, China), and The Native Antigen Company (Oxford, United Kingdom). In addition, the following defined strains were used in this study: *Streptococcus pneumoniae* 3 (ATCC 6303), *Streptococcus pneumoniae* 19A (ATCC 700674), *Escherichia coli* 41949 (Multiple O antigens:H26) (Crimson Biorepository), and *Escherichia coli* RS218 (NMEC O18:H7) (Kindly provided by James R. Johnson from the University of Minnesota). LPS from *Serratia marcescens* (L6136)*, Klebsiella pneumoniae* (L4268), *Salmonella enterica* serovar enteritidis (L6011), and LTA from *Enterococcus hirae* (L4015), *Staphylococcus aureus* (L2515), and *Streptococcus pyogenes* (L3140) were purchased through Sigma-Aldrich. *Mycobacterium tuberculosis* H37Rv components, which include lipoarabinomannan (LAM, NR-14848) and phosphatidylinositol mannoside 1,2 and 6 (PIM_1,2_, NR-14846 and PIM_6_, NR-14847), were obtained from BEI resources.

### Preparation of bacteria

Bacteria were subcultured in RPMI (Thermo Fisher Scientific, USA) 10mM glucose to a McFarland of 0.5 (equivalent to ~1e8 CFU/mL). Bacteria were grown to this logarithmic phase to ensure cell viability, and RPMI is used because it does not contain interfering MBL binding nutrients, such as yeast extract. The culture was then split - live bacteria were kept on ice while the other half were fragmented. Fragmented bacterial PAMPs were generated using antibiotics. Antibiotic treatment included the appropriate use of one of the following: cefepime (NDC 25021-121-20), ceftriaxone (NDC 60505-6104-4), meropenem (NDC 63323-507-20), amikacin (NDC 0703-9040-03), or vancomycin (NDC 0409-4332-49), at 1 mg/mL for ≥ 4 hours at 37°C 225 rpm. Testing by FcMBL ELLecSA was performed on titers of both live and fragmented bacteria at ≤ 1e7 CFU/mL. LPS endotoxin from Gram-negative bacteria was quantified using a limulus amebocyte lysate (LAL) assay ([Endosafe®] Charles River Laboratories, USA).

### Preparation of fungi, viruses, parasites, and bacterial cell wall antigens

Fungi species were primarily propagated in RPMI 10mM glucose, however other media, such as potato dextrose broth (Teknova, USA), were used to facilitate growth. In these cases, the fungal cells were pelleted at 3,000 × *g* for 5 minutes at 22°C (Eppendorf 5424, USA), washed 3x in 50mM Tris-HCl, 150mM NaCl, 0.05% Tween-20, 5mM CaCl_2_, pH 7.4 (TBST 5mM CaCl_2_) (Boston BioProducts, USA) to remove residual growth media, and then resuspended in TBST 5mM CaCl_2_. Testing by FcMBL ELLecSA was performed on titers of live fungi at ≤ 1e7 CFU/mL. Purified or inactivated viral, parasite, and bacterial cell wall antigens were resuspended or diluted in TBST 5mM CaCl_2_ for testing directly by FcMBL ELLecSA.

### FcMBL ELLecSA

The key metric used to quantify direct FcMBL binding to pathogen-associated molecular patterns (PAMPs) from bacteria, fungi, viruses, parasites, and bacterial cell wall antigens is a 96 well ELLecSA, which has been previously published [23]. The assay uses FcMBL coated superparamagnetic beads (1 µm MyOne Dynabead [Thermo Fisher Scientific, USA]) where FcMBL, biotinylated at the N termini of the Fc protein using an N-terminal amino-oxy reaction, is coupled to streptavidin beads in an oriented array (**Fig 1**). Each sample is screened using 5 µg of the FcMBL beads, 200 µL test sample, and 800 µL TBST 5mM CaCl_2_ supplemented with 10mM glucose (50mM heparin is added if testing blood). PAMPs in the test sample are captured by FcMBL for 20 minutes at 22°C 950 rpm in a plate shaker (Eppendorf, USA). Using an automated magnetic-handling system (KingFisher^TM^ Flex [not shown]) (Thermo Fisher Scientific, USA), captured PAMPs are washed two times using TBST 5mM CaCl_2_, and detected with human MBL (Sino Biological) linked to horseradish peroxidase (MBL-HRP). Non-specific MBL-HRP is removed by 4 washes in TBST 5mM CaCl_2_, and PAMPs are quantified with 1-step ultra tetramethylbenzidine (TMB) substrate (Thermo Fisher Scientific, USA). Finally, the reaction is quenched with 1N sulfuric acid and results are read at the optical density 450 nm wavelength. Quantification of bound PAMPs is determined using a standard curve generated using yeast mannan – a known target for MBL (1 ng/mL mannan = 1 PAMP unit) [38]. PAMP units are multiplied back by the dilution factor (×5) of the test sample volume to give PAMPs/mL. Previously, a receiver operating characteristic comparison was performed for a small pilot sepsis patient study in which each sepsis blood draw was analyzed versus non-infected controls to determine an optimal ELLecSA threshold of 0.45 PAMP units [23]. Therefore, in this study we define and report FcMBL binding to a sample as having ≥ 2.25 PAMPs/mL. To confirm specificity of FcMBL binding, a negative control (FcMBL null) was used alongside FcMBL in the ELLecSA. The FcMBL null was engineered by introducing two residue mutations, E347A and N349A, into aktFcMBL (GenBank accession: KJ710775.1) to remove functional binding of the CRD of MBL. FcMBL null was purified and used to coat beads in the same fashion as FcMBL described above for direct comparison. FcMBL null beads did not support any binding to yeast mannan.

### Scanning electron microscopy

For visualization of live and fragmented bacteria on FcMBL beads, bacteria were captured with 128 nm FcMBL beads (Ademtech, France), spun down onto 13mm coverslips and fixed with 2.5% glutaraldehyde in 0.1M sodium cacodylate buffer (Electron Microscopy Sciences, USA) for 1 hour. Cover slips were incubated in 1% osmium tetroxide in 0.1M sodium cacodylate (Electron Microscopy Sciences, USA) for 1 hour. Ascending grades of ethanol dehydrated the sample before being chemically dried with hexamethydisilazane (Electron Microscopy Sciences, USA). Samples were then placed in a desiccator overnight. Dried samples are mounted on aluminum stubs, sputter-coated with a thin layer of gold particles, and imaged using a Zeiss Supra55VP microscope.

## Acknowledgments

We thank Vasanth Chandrasekhar for assistance in making the FcMBL beads, Shanda Lightbown for SEM images of *S. pneumoniae* 3, and Seth Kroll for image processing.

